# Whole-genome analysis of noncoding genetic variations identifies multigranular regulatory element perturbations associated with Hirschsprung disease

**DOI:** 10.1101/2020.04.08.032045

**Authors:** Alexander Xi Fu, Kathy Nga-Chu Lui, Clara Sze-Man Tang, Ray Kit Ng, Frank Pui-Ling Lai, Sin-Ting Lau, Zhixin Li, Maria-Mercè Gracia-Barcelo, Pak-Chung Sham, Paul Kwong-Hang Tam, Elly Sau-Wai Ngan, Kevin Y. Yip

**Affiliations:** Department of Computer Science and Engineering, The Chinese University of Hong Kong, Hong Kong; Department of Surgery, Li Ka Shing Faculty of Medicine, The University of Hong Kong, Hong Kong; Dr. Li Dak-Sum Research Centre, The University of Hong Kong, Hong Kong; School of Biomedical Sciences, Li Ka Shing Faculty of Medicine, The University of Hong Kong, Hong Kong; Department of Psychiatry, Li Ka Shing Faculty of Medicine, The University of Hong Kong, Hong Kong; Centre for Genomic Sciences, Li Ka Shing Faculty of Medicine, The University of Hong Kong, Hong Kong; Hong Kong Bioinformatics Centre, The Chinese University of Hong Kong, Hong Kong; CUHK-BGI Innovation Institute of Trans-omics, The Chinese University of Hong Kong, Hong Kong; Hong Kong Institute of Diabetes and Obesity, The Chinese University of Hong Kong, Hong Kong

## Abstract

It is widely recognized that the missing heritability of many human diseases is partially due to noncoding genetic variants, but there are multiple challenges that hinder the identification of functional disease-associated noncoding variants. The number of noncoding variants can be many times of coding variants; many of them are not functional but in linkage disequilibrium with the functional ones; different variants can have epistatic effects; different variants can affect the same genes or pathways in different individuals, and some variants are related to each other not by affecting the same gene but by affecting the binding of the same upstream regulator. To overcome these difficulties, we propose a novel analysis framework that considers convergent impacts of different genetic variants on protein binding, which provides multi-granular information about disease-associated perturbations of regulatory elements, genes, and pathways. Applying it to our whole-genome sequencing data of 918 short-segment Hirschsprung disease patients and matched controls, we identify various novel genes not detected by standard single-variant and region-based tests, functionally centering on neural crest migration and development. Our framework also identifies upstream regulators whose binding is influenced by the noncoding variants. Using human neural crest cells, we confirm cell-stage-specific regulatory roles three top novel regulatory elements on our list, respectively in the *RET, RASGEF1A* and *PIK3C2B* loci. In the *PIK3C2B* regulatory element, we further show that a noncoding variant found only in the affects the binding of the gliogenesis regulator NFIA, with a corresponding down-regulation of multiple genes in the same topologically associating domain.

## Introduction

Hirschsprung (HSCR) disease is a rare, complex genetic disease characterized by missing enteric ganglia in various portions of the hindgut^1^. It is caused by failed migration, proliferation, differentiation, or colonization of enteric neural crest (NC) cells, which disrupts enteric nervous system (ENS) development^2,3^. Phenotypic severity of the disease is determined by the length of colonic aganglionosis and can be classified into short-segment HSCR (S-HSCR; 80% of cases), long-segment HSCR (L-HSCR; 15-20% of cases) and total colonic aganglionosis (TCA; up to 5% of cases)^3,4^.

HSCR is long recognized to be highly heritable (80%-97% for S-HSCR and ∼100% for L-HSCR), while around 80% of HSCR cases are sporadic (>95% for S-HSCR)^2,5^. The incidence of HSCR varies across ethnic groups, from 1.4 to 2.8 in every 10,000 new births, with the highest incidence rate in Asia^1,3,6^. The genetic etiology of HSCR is multifactorial, involving rare and common, coding and noncoding variants in genes playing different roles in ENS development. Among the different subtypes of HSCR, S-HSCR is genetically most complex – Whereas L-HSCR and TCA cases are mostly autosomal dominant, the less severe S-HSCR subtype follows a complex, non-Mendelian inheritance pattern, and has a male-to-female ratio of 4:1 (Refs. 2,7) with the reason behind it not fully known.

Damaging rare variants at protein-coding sequences associated with HSCR have been found in many genes involved in ENS-development. Among them, *RET*, encoding a receptor tyrosine kinase, is responsible for around 50% of familial cases and 10-20% of sporadic cases^3^. Other genes previously reported include *BACE2, DNMT3B, ECE1, EDN3, EDNRB, FAT3, GDNF, GFRA1, KIAA1279, L1CAM, NRG1, NRG3, NRTN, NTF3, NTRK3, PHOX2B, PROK1, PROKR1, PROKR2, PSPN, SEMA3A/C/D, SOX10, TCF4*, and *ZFHX1B* ^8–13^, which together account for around 5% of all cases. These rare damaging variants often have incomplete penetrance and explain only a small portion of the phenotypic variance.

In contrast, the most well-known common variant, rs2435357, located in intron 1 of *RET* and affects SOX10 binding to a *RET* enhancer, explains 10-20 times of the phenotypic variance as compared to the rare damaging *RET* variants^7,14,15^. Recently, some additional noncoding variants have also been found associated with HSCR. For example, two other variants in *RET* enhancers act synergistically with rs2435357 on HSCR risk^15^, while variants in four noncoding elements around *RET* and *SEMA3* were found in 48.4% of cases and 17.1% of controls, with the presence of five or more variants in these regions corresponding to a high disease risk^11^. However, despite the ever-growing knowledge about HSCR, there is still a considerable portion of cases that cannot be explained by the cataloged coding or noncoding variants.

The emerging studies of HSCR-associated common variants in *RET* enhancers and other functional noncoding regions^11,14,15^ and the abundance of noncoding signals from genome-wide association studies of many human traits in general^16^ both suggest that noncoding regions would be a valuable territory to explore in the quest of explaining the missing heritability in HSCR.

Existing methods for studying noncoding genetic variants can be broadly classified into site-based and region-based methods. Site-based methods (reviewed in Ref. 17) aim at prioritizing the variants according to their potential functional effects, based on information such as evolutionary conservation, sequence patterns and epigenomic signals. Such information has been recorded in various annotation databases^18–21^, including cell/tissue-specific information indicative of the functional potential of noncoding regions including chromatin accessibility, histone modifications and transcriptional activities^22–25^. In contrast, region-based methods consider genomic regions of potential functional significance as the basic units, such as enhancers^26^, promoters^27,28^, contact regions in three-dimensional genome architecture^29,30^, or combinatorial categories^27^. The functional potential of these regions is usually quantified by either the frequency of genetic variants (burden test and its derivatives) or more complex measures, involving sequence kernel association test^31^, convolutional neural network (CNN) kernels that resemble transcription factor (TF) binding site motifs^32–34^, or gene expression models^33,35,36^.

In general, both site-based and region-based methods face several major challenges, namely 1) there is a large number of sites/regions to consider genome-wide, leading to low statistical power due to multiple hypothesis testing correction, 2) many genetic variants at noncoding regions do not have direct functional effects, including those in linkage disequilibrium with the actual functional variants, 3) epistatic interactions can exist among genetic variants, making it unsuitable to consider each variant separately^37^, 4) genetic variants at different regulatory regions of the same gene/pathway may result in the same convergent effect, and therefore each of them may not have high recurrence in the study subjects, and 5) the loci of different genetic variants could be related to each other not by their genomic locations but by the upstream regulators that commonly bind them, which are more non-trivial to determine.

In this study, we propose a new noncoding variant prioritizing framework, Multigranular Analysis of Regulatory Variants on the Epigenomic Landscape (MARVEL), that addresses these five challenges. Applying our framework to the whole-genome sequencing (WGS) data generated from an S-HSCR cohort (431 cases and 487 ethnically matched controls), we identify disease-associated enhancers, promoters, and genes based on epigenomic data generated from the human pluripotent stem cell (hPSC)-derived enteric NC-like cells (hNC). This is by far the largest WGS data set of S-HSCR with both cases and controls coming from a single population. By aggregating lower-level (genomic variants and TF binding motifs) information, marginally significant signals emerge as more significant associations at a higher level (enhancers, promoters, and genes) that converge to specific functional pathways and upstream regulators.

## Results

### A novel association framework for the noncoding genome

MARVEL is designed to address the five challenges listed above (Figure 1, Methods). First, we use cell-type-specific epigenomic data to define a target set of active regulatory elements in the relevant cell types, in order to limit the number of statistical tests to be performed (Figure 1a). Second, in these target elements, we use sequence motifs as a proxy to evaluate the functional potential of the genetic variants^38^, which helps prioritize the genetic variants and their corresponding regulatory elements among those in linkage disequilibrium (Figure 1b). Third, for each target element, we consider all genetic variants simultaneously to reconstruct the sample-specific sequence for evaluating their joint functional effects (Figure 1c). For instance, if two genetic variants have opposite effects on the binding strength of a TF according to its sequence motif, the net expected effect would be smaller than the sum of their individual absolute effects. Fourth, our framework considers multiple occurrences of a motif in the same regulatory element (Figure 1d) or different regulatory elements of the same gene (Figure 1e) in a joint manner. As a result, a gene would be considered significantly perturbed when its regulatory elements have frequent motif-disrupting genetic variants even if in different subjects these regulatory elements contain different sets of genetic variants^15,39^. The joint effect of multiple motifs is further captured by first selecting an important combination of the motifs using a feature selection procedure (Figure 1f) followed by an association test (Figure 1g), which together identify motifs whose match scores in these regulatory elements are significantly associated with the target phenotype, adjusted for covariates such as age and sex of the subjects. The identified phenotype-associated regulatory elements and genes are further investigated by additional analyses such as gene set enrichment and single-cell expression analyses (Figure 1h). Fifth, by using sequence motifs as functional proxy, we also identify the potential upstream regulators whose binding to different target elements are frequently perturbed, which can be either loss or gain of binding. By identifying regulators whose binding motifs are commonly perturbed in the regulatory elements of different genes (Figure 1i), we identify both the core regulators involved in the phenotype^40^ and genes potentially involved in the same pathways.

**Figure 1.**
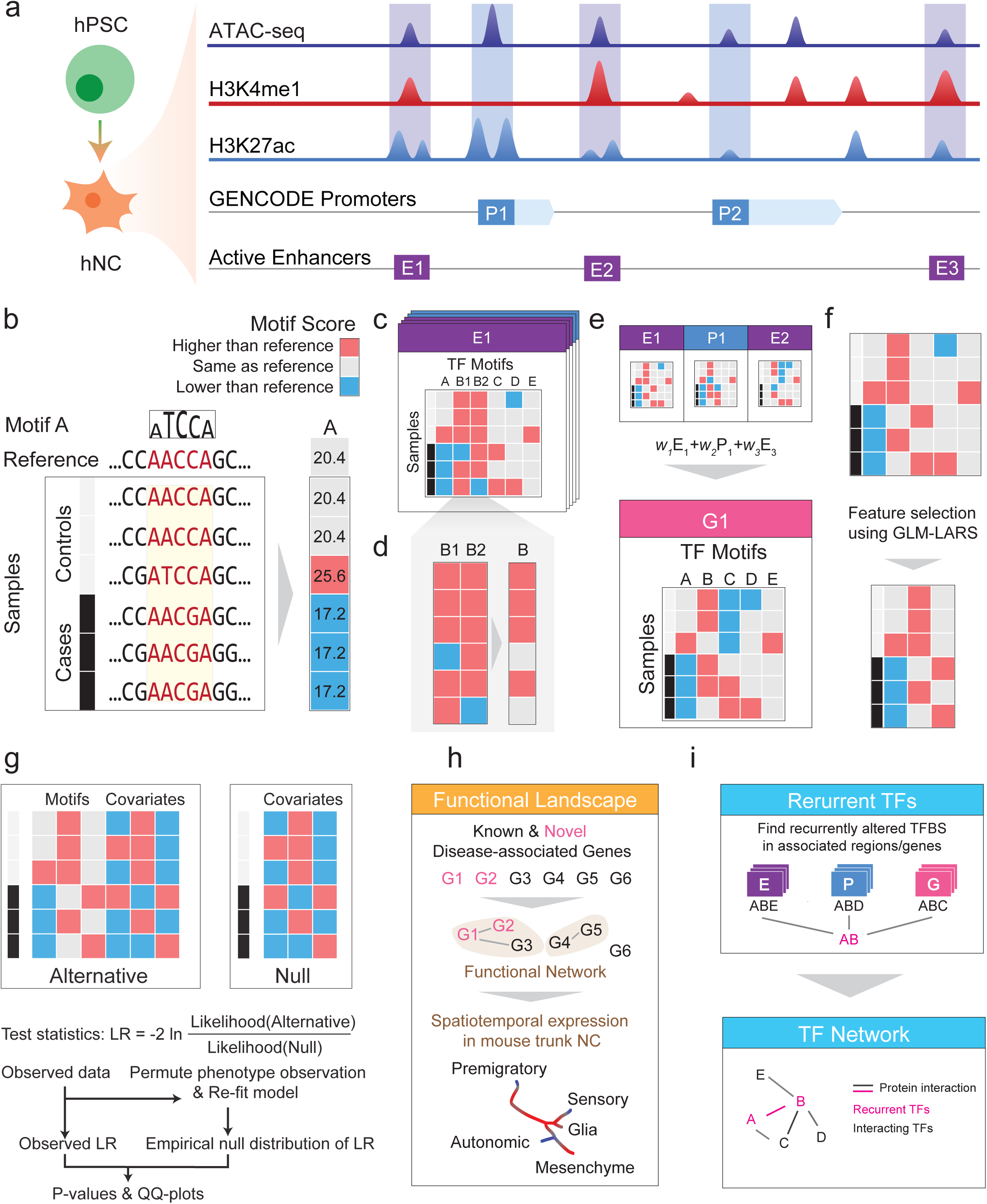
Schematic overview of MARVEL. (**a**) Epigenomic data of relevant cell type (hNC in the case of HSCR) are integrated with a gene annotation set to identify the active regulatory elements relevant to the phenotype of interest. (**b**) In each regulatory element, the functional significance of genetic variants is evaluated by their perturbation to TF sequence motifs. (**c**) Since the perturbation effects of multiple genetic variants may not add up linearly, they are considered together to reconstruct the sample-specific sequences, based on which the overall change of TF motif match scores is determined. (**d**) For motifs with multiple appearances within the same regulatory element, their match scores are aggregated to give a single score. (**e**) At a higher level, if a gene involves multiple regulatory elements, the aggregated match scores of a motif in the different elements can be further aggregated into a single score. This is done in the gene-based analysis. (**f**-**g**) The aggregated match score matrix of all the motifs for a regulatory element/gene is used as the input of an association test, which selects a subset of the most informative motif features (f) and compares a model involving both these selected features and the covariates with a null model that involves only the covariates using likelihood ratio (LR) test (g). (**h**) The regulatory elements and genes identified to be significantly associated with the phenotype can be further studied by other downstream analyses, such as gene set enrichment and single-cell expression analyses. (**i**) TFs with recurrently perturbed match scores in different regulatory elements are collected to infer a network that highlights the phenotype-associated perturbations.

Using simulated data (Methods), we verified that MARVEL is able to select informative motifs truly associated with the phenotype, including motifs whose match scores are affected by genetic variants at different allele frequencies, effect sizes, and correlations among each other (Supplementary Figure 1). We also verified that the P-value and effect size distributions produced by MARVEL clearly separated the truly disease-associated regions from the background (Supplementary Figure 2).

### Novel noncoding regulatory elements and gene loci associated with S-HSCR

We applied MARVEL to analyze our S-HSCR WGS data (Methods). Briefly, target enhancers and promoters were defined using chromatin accessibility and histone modification data from hNC, which were respectively used in an enhancer-based and a promoter-based analysis. A gene-based analysis was also performed by considering both promoters and enhancers potentially regulating different transcript isoforms of each gene, with the motif scores of different regulatory elements weighted according to their expected chromatin contact frequencies with the transcription start site (TSS) in hPSCs. This gene-based analysis allowed the detection of genes with frequent perturbations by multiple low-recurrence genetic variants at different regulatory elements. Association tests were then performed based on the motif scores in each of these three sets of target regions in turn, with 150,828 enhancers, 87,461 promoters, and 24,557 genes, respectively.

We defined all regions above the 0.95 confidence interval of the null in the quantile-quantile plot as loosely associated with S-HSCR, and all regions passing the FDR threshold of 0.1 as significantly associated (Figure 2a, Supplementary Table 1). Quantified by the area under the receiver-operator characteristic (AUROC, Methods), the loosely associated regions had generally larger effect sizes than random regions in the same sets of target regions (Figure 2b). Comparing the three sets of target regions, there were more enhancers and genes significantly associated with S-HSCR as compared to promoters. Noncoding variant association methods that focus on the promoter regions would have missed the many associated enhancers and aggregated regulatory regions of the genes. As a negative control, we repeated the procedures on a background set of enhancers that do not overlap the hNC enhancers (Methods), and found none of them loosely associated with S-HSCR (Supplementary Figure 3).

**Figure 2.**
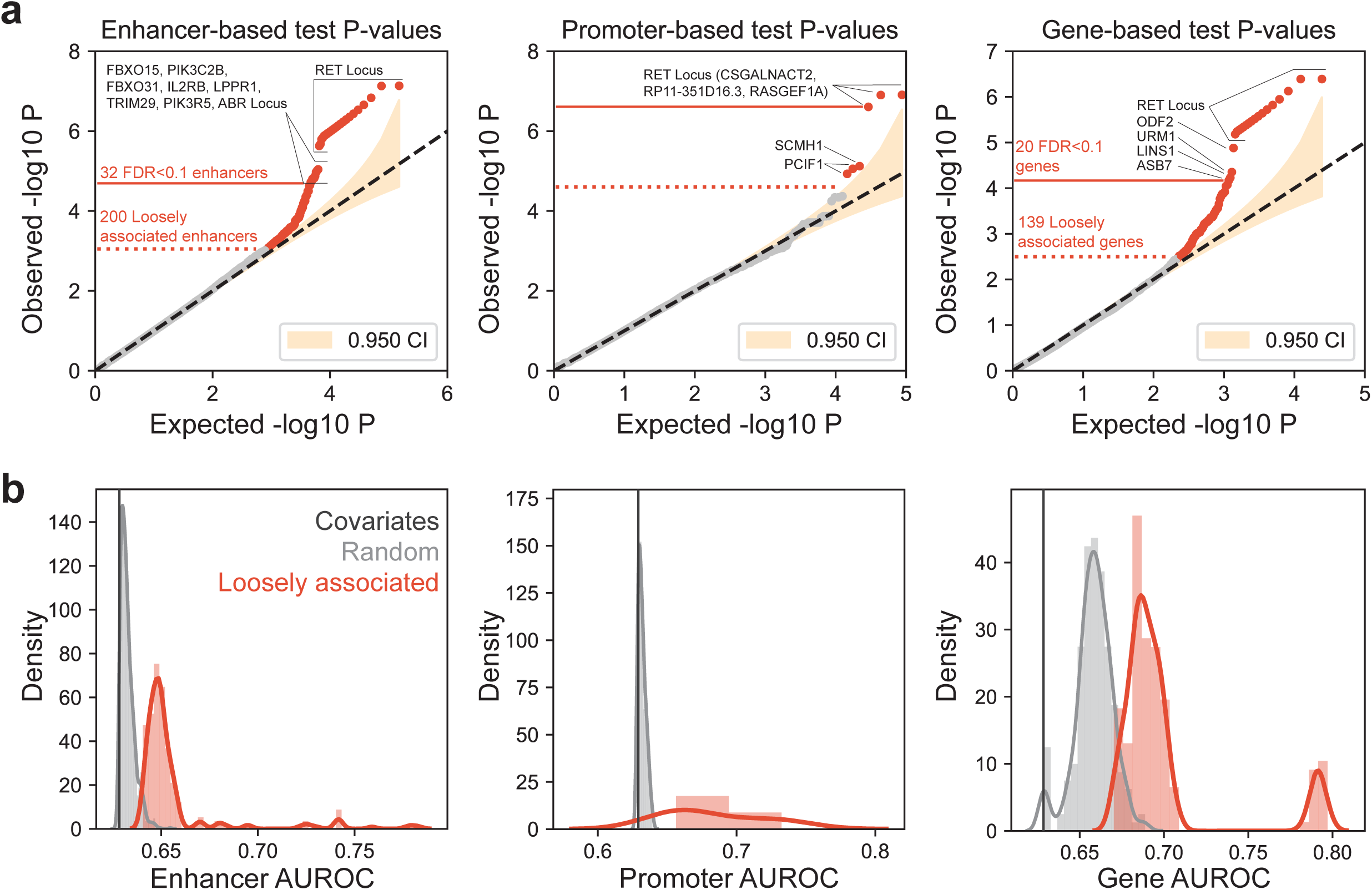
Association test results. (**a**) QQ-plots of association P-values in the enhancer-based, promoter-based, and gene-based tests. In each plot, the yellow shaded area shows the 95% confidence interval according to beta error distribution. The dotted red line marks the threshold for the loosely associated regions, above which all the regions are outside the 95% confidence interval. The solid red line marks the threshold for significantly associated regions, above which all regions have an FDR Q-value<0.1. The significantly associated regions are a subset of the loosely associated regions. (**b**) Comparison of the AUROC value distributions of the loosely associated regions with random regions.

**Table 1.** Significant S-HSCR associated enhancers, promoters and genes. In all three cases, regions near *RET* are collapsed into one entry. For the enhancer list, the nearest genes of each enhancer were identified based on the distance between the center position of the enhancer and the TSSs of genes. Expressed and unexpressed TFs are defined based on our hNC RNA-seq data, with a cutoff threshold of FPKM (fragments per kilobase of exons per million reads)=1. *: The smallest P-values that could be obtained from our permutation tests

| Region | P-value(s) | Altered motifs of TFs expressed in hNC |  |
| --- | --- | --- | --- |
| Direction of binding change associated with increased disease risk: |  | Gain | Loss |
| **Significantly associated enhancers (and 3 corresponding nearest genes of each)** |  |  |  |
| 23 enhancers near *RET* | 7.42E-08* to 2.35E-06 |  |  |
| chr18:73918014-73919015<br>(*FBXO15*, *TIMM21*, *CYB5A*) | 9.10E-06 | CPEB1, NFATC1, NFIC, ZNF322 | LEF1, SP3, TBX1, WT1 |
| chr1:204456576-204457577<br>(*PIK3C2B*, *PPP1R15B*, *MDM4*) | 9.77E-06 | EGR2, HOXC6, TWIST1, ZFP82, ZNF528 | EGR1, HAND1, NFIA, ZNF816 |
| chr16:86954235-86955236<br>(*C16orf95*, *FOXL1*, *FBXO31*) | 1.06E-05 | ETS2, NFAT5, PITX2 | FLI1, SP2, TAF1, ZEB1, ZNF317, ZNF418 |
| chr22:37128477-37129478<br>(*IL2RB*, *TMPRSS6*, *C1QTNF6*) | 1.33E-05 | EGR2, HAND1, KLF16, ZNF263 | NHLH1, ZNF322 |
| chr9:100937755-100938756<br>(*LPPR1*, *MURC*, *BAAT*) | 1.51E-05 | FOXH1, TBX15, ZNF816 | AR, ETV7, TGIF2LX, ZNF335 |
| chr11:120020704-120021705<br>(*TRIM29*, *OAF*, *POU2F3*) | 1.59E-05 | ARID5B, SOX9, TGIF1, ZNF816 | IRF2, NKX6-1, TEAD2, ZNF394 |
| chr17:8915658-8916659<br>(*PIK3R5*, *PIK3R6*, *NTN1*) | 1.79E-05 | NR2F1, POU2F2, SP3, UBP1, ZNF148 | ETV3, NR2F6, ZNF317 |
| chr17:958411-959412<br>(*NXN*, *TIMM22*, *ABR*) | 1.9E-05 | EVX1, FOXG1, PITX2, ZNF770 | FOXD2, PAX5, ZNF667 |
| **Significantly associated promoters** |  |  |  |
| 3 promoters near *RET* | 1.24E-07* to 2.48E-07 |  |  |
| **Significantly associated genes** |  |  |  |
| 16 genes near *RET* | 4.07E-07* to 6.05E-06 |  |  |
| *ODF2*<br>(chr9:128455155-128501292) | 9.27E-06 | E2F4, MYOD1, ZBTB4, ZNF274 | NR1I3, SMAD3, TGIF2, ZFP64 |
| *URM1*<br>(chr9:128371319-128392016) | 4.35E-05 | E2F4, ELK3, MYOD1, ZBTB4, ZNF274 | NR1I3, SMAD3, ZFP64 |
| *LINS1*<br>(chr15:100559369-100603230) | 5.89E-05 | FOXC1, FOXF1, ZNF394 | HAND1, MECP2, MTF1, ZSCAN31 |
| *ASB7*<br>(chr15:100602534-100651705) | 6.81E-05 | FOXC1, FOXF1, ZNF394 | HAND1, MECP2, MTF1, ZSCAN31 |

As expected, all three analyses identified various regions near the *RET* locus as significantly associated with S-HSCR (Table 1). The enhancer-based tests identified 30 significant enhancers, including 23 enhancers near *RET* (within 200kbp from the *RET* TSS). The top S-HSCR-associated enhancer (chr10:43086011-43087012) contains the well-known HSCR-associated common SNP rs2435357. Another RET-locus enhancer (chr10:43064374-43065375) has VDR binding loss more frequently in cases than in controls (odds ratio: 7.93, 95%CI: 5.88 to 10.70, P<0.0001). Interestingly, VDR is a vitamin-D receptor that been shown to directly regulate RET expression^41^. The promoter-based tests identified only three significant promoters, all of which are promoters of immediate neighboring genes of *RET* (*CSGALNACT2, RASGEF1A* and *RP11-351D16*.*3*). The gene-based tests identified 20 significant genes, including *RET* itself and 15 genes near it (within 700kbp from the *RET* TSS).

More interestingly, we also identified eight enhancers and four genes significantly associated with S-HSCR that are not close to *RET* (Table 1). Among the eight enhancers, the one on chromosome 1 (chr1:204456576-204457577) overlaps intron 10 of *PIK3C2B*. PIK3C2B is a phosphoinositide 3-kinase (PI3K) family protein, which is believed to have important roles in the signal integration and transduction in hNC^42^. Binding sites of TWIST1 and NFIA are altered in this enhancer, both with increased match scores in 6 cases but not in any controls (Table 1). TWIST is an hNC specifier^43^ while NFIA has been shown to regulate gliogenesis, which is also crucial to a functional ENS^44–46^. Interestingly, another enhancer on chromosome 17 (chr17:8915658-8916659) is also close to 2 other PI3K family genes, *PIK3R5* and *PIK3R6*. The two enhancers on chromosomes 18 and 16 (chr18:73918014-73919015 and chr16:86954235-86955236) are within/close to F-box genes (*FBXO15* and *FBXO31*, respectively). F-box proteins bind to CUL1 to form SCF (SKP-CUL1-F-box protein) E3 Ubiquitin Ligase complexes, which mediate ubiquitination of proteins that regulate the cell cycle and diverse neuronal activities^42,47^. In particular, FBXO31 has been shown to regulate neural morphogenesis and migration^48^. The genes closest to the enhancer on chromosome 22 (chr22:37128477-37129478) are not clearly related to HSCR, but *SOX10* and *CDC42EP1* are within 100kbp from it. *SOX10* is an HSCR-associated gene^49–51^, while CDC42EP1, a CDC42 binding protein, is involved in neural crest cell directional migration by regulating the actin polymerization^52^. Another enhancer on chromosome 17 (chr17:958411-959412) is also potentially related to CDC42 since its neighboring gene, *ABR*, encodes a protein that is able to promote Rho activity and reduce CDC42 activity^53^, both of which are important for actin regulation and cell polarization during migration^52,53^. The enhancer on chromosome 9 (chr9:100937755-100938756) is close to *LPPR1*, which encodes a member of the plasticity-related gene (PRG) family. Members of the PRG family mediate lipid phosphate phosphatase activity in neurons and are known to be involved in neuronal plasticity^54,55^, potentially also involved in ENS development^56^. The enhancer on chromosome 11 (chr11:120020704-120021705) is close to *TRIM29*, a gene that regulates epithelial-to-mesenchymal transition (EMT), which is an important process shared by both NC migration and cancer metastasis^57^.

We compared these enhancer-based results with three commonly used single-variant and region-based association tests based on the variants in the hNC enhancers (Methods). The single-variant Wald test identified 69 variants at FDR<0.1 and in total 160 variants above the 0.95 confidence interval of the null in the quantile-quantile plot (Supplementary Figure 4a). Among these 160 variants, 69 of them were located within 31 of the loosely associated enhancers identified by MARVEL, leaving the remaining 169 loosely associated enhancers identified by MARVEL not detected by this single-variant test, among which are 11 of the significantly associated enhancers including all the 8 enhancers outside the *RET* locus (Supplementary Figure 4b). Among the 91 loosely associated variants uniquely identified by the Wald test, 45 of them did not overlap with any sequence motifs. Furthermore, 20 of these 45 variants are in linkage disequilibrium with some loosely associated variants that overlapped sequence motifs, suggesting that they may not be functional themselves. As for the region-based association tests CMC and SKAT-O, none of the hNC enhancers were found to be either significant or loosely associated (Supplementary Figure 4c-d). These results show that MARVEL can identify S-HSCR associated regions missed by these commonly used methods.

The four significant genes not close to the *RET* locus come from two loci respectively on chromosomes 9 and 15, with no loosely associated enhancers or promoters near them. Each of these genes was thus found to be significantly associated with S-HSCR due to weak association signals that distribute across multiple regulatory regions of it. Indeed, a significantly larger fraction of enhancers within 1Mbp from the TSSs of these genes have P-values smaller than 0.1 as compared to the set of all enhancers (P=0.025, Fisher’s exact test). One of the genes in these loci is *ODF2*, which encodes the Cenexin protein. Previous studies have demonstrated that in various mammalian cell lines, Cenexin controls centrosome positioning during directional cell migration^58^. Since centrosome acts as a steering machine for cell movement and stabilizes the migration direction^58,59^, ODF2 might help stabilize the direction of NC migration.

### Loosely associated signals aggregate into functional pathways

Having found that genetic variants weakly associated with S-HSCR at different regulatory regions aggregated to give stronger signals at the gene level, we next investigated whether the association signals could be further aggregated into functional pathways. We performed this investigation by taking the 552 genes loosely associated with S-HSCR from our enhancer-based, promoter-based and gene-based analyses (Methods), and connected those of them with functional interactions in Reactome^60^. A total of 114 genes were connected, by 316 functional interactions. Both values were significantly larger than those of random gene sets of the same size (P=0.018 for the number of connected genes and P=0.031 for the number of interactions), suggesting that these S-HSCR genes were indeed functionally related.

To explore the actual functions of these S-HSCR genes, we expanded this gene set by adding known HSCR genes and genes important in ENS development or NC migration (Methods), and found that the resulting genes formed five main clusters according to their functional interactions (Figure 3a). These five clusters are respectively related to (1) chemotaxis and cell-cell signaling; (2) cell adhesion, migration, and interaction with extracellular matrix; (3) PI3K/PKC/MAPK/PPARG signaling; (4) E3 Ubiquitin Ligase complexes; and (5) transcriptional regulatory factors. They are highly related to NC migration and its development (Figure 3b) and involve many genes loosely associated with S-HSCR (Supplementary Table 2).

**Figure 3.**
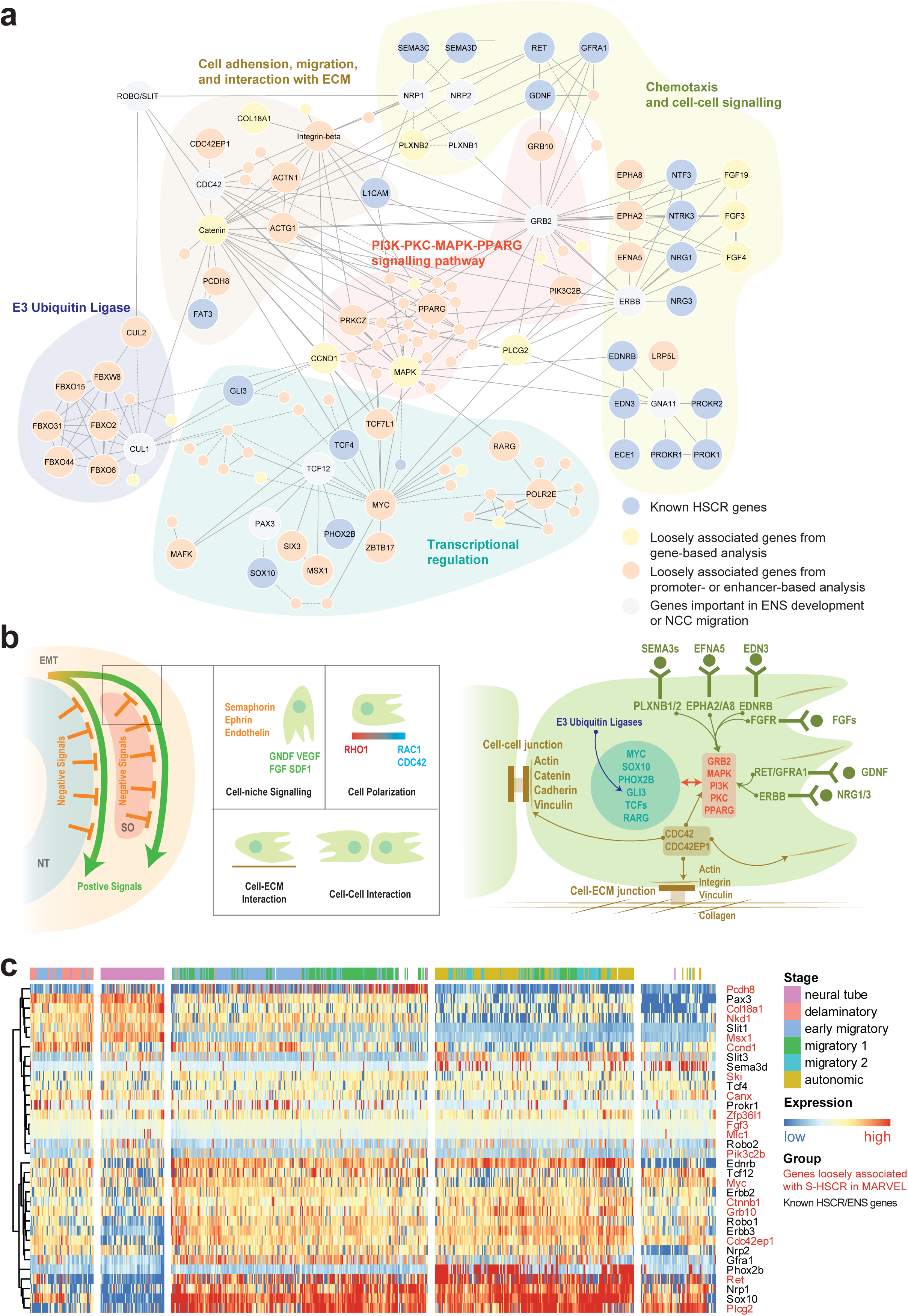
Functional landscape of S-HSCR associated genes. (**a**) Functional interactions among genes loosely associated with S-HSCR, known HSCR genes and genes important in ENS functions or NC migration. Each node corresponds to a gene and each edge corresponds to a functional interaction cataloged in Reactome. Genes of particular interest are shown in bigger nodes, labeled with their names. (**b**) Schematic illustration of some biological processes and genes involved in NC migration^61^. Colors of gene names follow their functional categories in Panel a. Abbreviations: NT, neural tube; SO, somites. (**c**) Spatialtemporal expression profiles of mouse trunk NCs. Each row corresponds to the stage-specific expression pattern of a mouse homolog of a human gene shown in Panel a. Genes identified by MARVEL as loosely associated with S-HSCR are shown in red. Each column corresponds to a single cell with the predicted stage label taken from the original publication^70^. Both the rows and the coumns were clustered using hierarchical clustering with Pearson correlation as the similarity measure, and the columns are divided into 5 partitions according to the clustering results.

Chemotaxis signaling is used to direct NC to target tissue (positive signaling) and works against the negative signals from the neural tube or somite to promote NC delamination and migration^61^. Additionally, cell polarization is critical during the directional migration of NC. Under the regulation of the RAC1-RHO1 gradient, actin is selectively polymerized on one side of the cell, extending the cell along the migration direction^52,61^. The extracellular matrix (ECM) serves as the road of NC migration^61^ and collagens are basic structures in the ECM required for successful NC migration^62^. Cell-ECM interactions such as focal adhesion through Integrins and Vinculins provide both structural support and signaling sensor for migrating NC. In a previous study, mutations of Vinculin in HSCR patients were shown to affect its function in focal adhesion^63^. Cell-cell adhesion through Catenin, Cadherin and Vinculins is important for collective NC migration^61,64^.

Signals from cell-ECM interactions, cell-cell interactions and chemotaxis are integrated inside NC through several signaling pathways including PI3K, MAPK/ERK, PPAR-γ and PKC pathways^65^. Through these pathways, signals from the environment are transformed into regulatory signals, controlling the activation and deactivation of genes.

An additional layer of regulation of these cellular signals was found at protein level. For instance, E3 ubiquitin ligase is required by GLI3, a known HSCR-associated TF, to switch between its activator and repressor forms^66^ by which GLI regulates the differentiation and patterning of enteric NC^67^.

Besides known HSCR TFs, we also found several novel TFs loosely associated with S-HSCR such as MYC and ZBTB17 (Supplementary Table 2) that are known to regulate the size of the pool of premigratory NC^68^ and RARG that is important in retinoic acid signaling pathway, a pathway crucial for enteric NC migration^69^.

Overall, our results reveal an unprecedentedly broad linkage between noncoding genetic variants associated with S-HSCR and NC genes, and clearly demonstrate that these genetic variants distribute across many different regulatory regions and genes that converge to key functional pathways.

As an additional way to explore the functions of the genes loosely associated with S-HSCR, we examined the spatiotemporal expression of their mouse homologs using a trunk NC single-cell RNA-seq (scRNA-seq) data set^70^ (Methods). Some of the known HSCR genes and loosely associated genes have stage-specific expression (Figure 3c). Based on the clustering result, these genes can be roughly divided into two groups, namely genes with higher expression in neural tube as compared to other stages (mainly in the upper cluster in Figure 3c), and genes with higher expression in the migratory stage and autonomic neuron stage (mainly in the lower cluster).

In the higher cluster, *Col18a1*, of which the human homolog was identified as loosely associated with S-HSCR in our gene-based analysis, has an expression profile similar to *Pax3*, a known regulator of Ret^71^. Interestingly, a previous study proposed that Col18a1 is secreted by NCs to regulate their migration during ENS development^62^. Another interesting case is *Pcdh8*, identified from our enhancer-based analysis, which encodes a Protocadherin and has specifically higher expression in the “migration 2” stage. The *Xenopus* ortholog of *Pcdh8, PAPC*, has been shown to play important roles in cell adhesion and NC migration^72^.

The lower cluster contain two well-known HSCR genes, *Ret and Phox2b. Plcg2*, identified in our gene-based analysis, has a similar expression pattern with another known HSCR gene, *Sox10*. The human ortholog of *Plcg2* was previously proposed to be a potential candidate of HSCR^73^. In addition to these examples, several other known ENS genes (*Ednrb, Robo1, Erbb3, Nrp2*) and genes loosely associated with S-HSCR (*Myc, Ctnnb1, Grb10, Cdc42ep1)* also have similar expression patterns in the marked stages. Some of these loosely associated genes have been shown to play important roles (Supplementary Table 2) in the pathways in Figure 3a.

### Upstream regulators of S-HSCR associated genes

One major advantage of our framework is its ability to identify not only the disease-associated regulatory elements or genes but also their upstream transcriptional regulators whose binding sites are altered by the genetic variants. Given the large number of enhancers loosely associated with S-HSCR found in our enhancer-based analysis, we developed a permutation test to identify TF binding motifs whose match scores are significantly more altered in these enhancers than the background set of all enhancers (Methods). This analysis identified 48 TFs that we will refer to as “recurrent TFs” (Supplementary Table 3). Interestingly, among the top 100 promoters with the strongest association P-values with S-HSCR, 73 of them also had at least one of these recurrent TFs’ motifs selected by GLM-LARS as a key motif, suggesting that in NCs these recurrent TFs regulate gene expression through both promoters and enhancers.

The recurrent TFs form an extensive network by way of physical protein-protein interactions (PPIs), with MYC and SMAD proteins acting as interaction hubs (Figure 4a). Interestingly, the premigratory NC pool size regulator ZBTB17-MYC and the retinoic acid receptor RARG discussed above in the functional pathway analysis are also contained in this network, showing that both the expression of these genes themselves and their downstream targets could be perturbed in S-HSCR. Some other interactions in this network are also known to have important functional roles. For example, Myc interacts with Smad to activate expression of Snails, which are essential TFs that regulate EMT of NC^68,74^. The E2f proteins have been shown to regulate neuronal migration^75^. There are also reported HSCR genes in the network, such as GLI3^67^.

**Figure 4.**
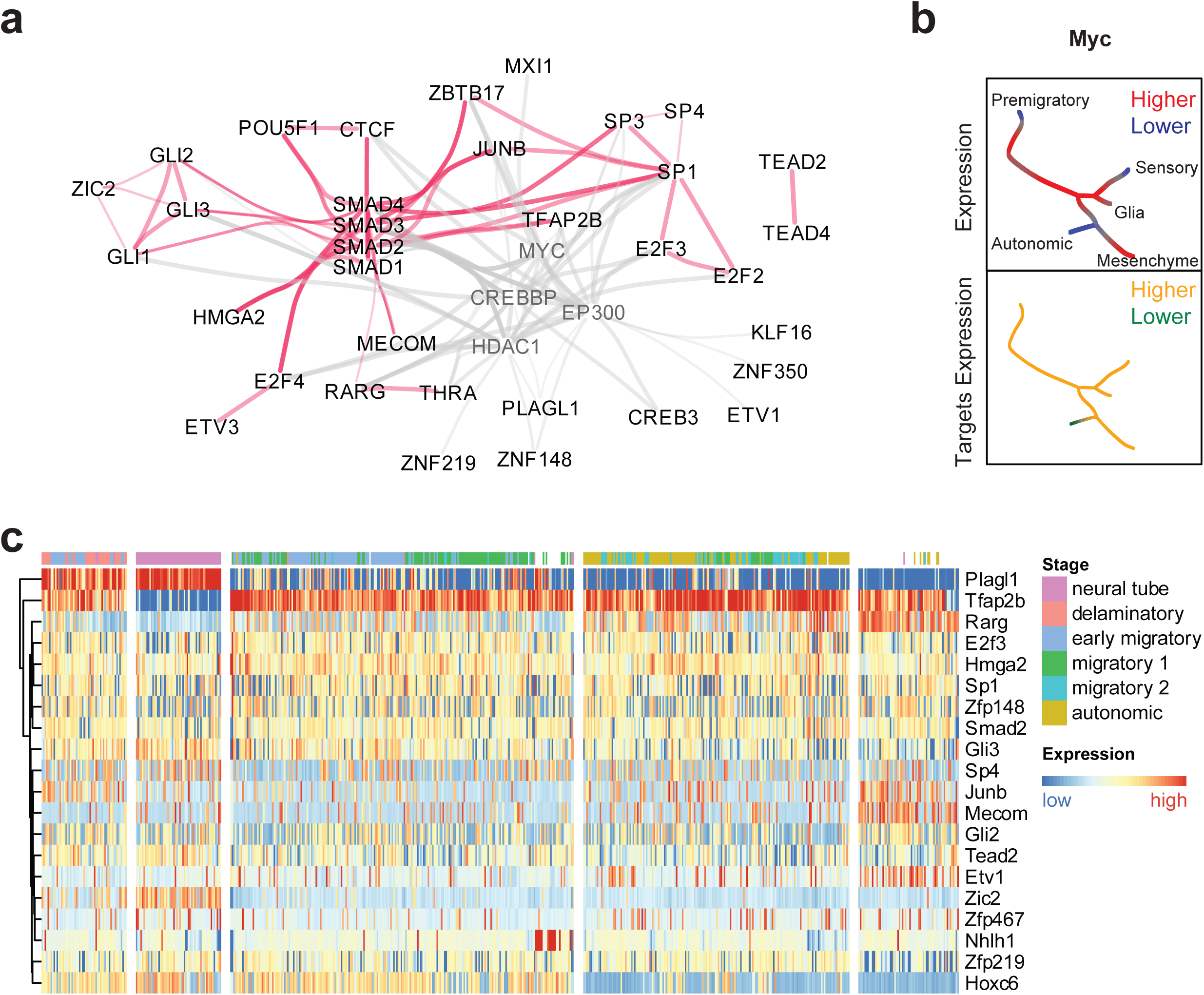
Analysis of the recurrent TFs. (**a**) PPIs among the recurrent TFs (black) and several other proteins frequently interacting with them (gray). Direct interactions among the recurrent TFs are shown in red, while direct interactions involving the other proteins are shown in gray. Recurrent TFs that have no interactions with other proteins in this figure are excluded. (**b**-**c**) Spatiotemporal expression of Myc and its potential regulated genes (b) and recurrent TFs with stage-specific expression profiles (c) in mouse trunk NCs.

Using the same mouse trunk NC scRNA-seq data described above, we found that the mouse homologs of some of the recurrent TFs have stage-specific expression. For example, *Myc* is down-regulated in premigratory, autonomic, and sensory neuron branches as compared to other cell stages. Correspondingly, the average expression of genes potentially regulated by Myc (from Ref. ^70^) is also specifically lower in the autonomic branch (Figure 4b). The recurrent TFs with the strongest across-cell expression variance are *Tfap2b* and *Plagl1* (Figure 4c). *TFAP2B* (human homolog of *Tfap2b*) is a known migratory NC marker^76^. *Plagl1* is upregulated in neural tube and in the delaminatory stage, which potentially regulates the composition of the ECM^77^ and thus potentially affects the migration environment of NC. One gene that has a similar expression pattern as *Plagl1* is *Zic2, which* regulates the production of NC^78^.

### Genetic variations at enhancers significantly associated with S-HSCR confer cell stage-specific expression of *RET*

To delineate the potential biological roles of our newly identified S-HSCR associated enhancers in disease susceptibility and pathogenesis, we employed hPSCs to model the development of the human ENS. We started with the most well-characterized HSCR-associated common SNP, rs2435357, where the risk allele *T* decreases the expression of *RET* and contributes to disease prevalence^79^. As mentioned above, this SNP resides in an enhancer within intron 1 of *RET* that was found to be significantly associated with S-HSCR in our enhancer-based analysis (Table 1). It overlaps an hNC-specific ATAC-seq peak (Figure 5a), suggesting that it may be implicated in the cell stage-specific expression of *RET*. To test this hypothesis, we introduced the risk allele *T* into a control hPSC line using a CRISPR/Cas9-mediated homology-directed repair (HDR) system with a specific single guide RNA (sgRNA) targeting the rs2435357(*C*/*C*) locus and single-stranded oligonucleotides (ssODNs)^80–82^ containing the *T* allele (Figure 5b). The control and mutant (UE-rs2435357) hPSC lines were then used to generate neural crest (hNC: SOX10^+^) and were subsequently directed to the neuronal lineage to make the early enteric neuronal progenitors (hNP: TUJ1^+^, *ELAVL4*^*+*^), recapitulating the two key developmental stages during the formation of the ENS as previously described^63,83^ (Figure 5c). The single base (*C*>*T*) conversion alone did not affect hNC induction and the derivation of hNPs from hPSCs. Intriguingly, we found that *RET* is highly expressed in the neuronal lineage committed hNCs (hNPs) and hPSCs, while its expression remains at a low level before committing to neuronal lineage, suggesting that the cell stage-specific enhancers are likely involved to mediate the dynamic expression of *RET* during the development of the ENS. In concordant with this observation, the *C*>*T* conversion in rs2435357 significantly reduced *RET* expression in hNPs derived from the UE-rs2435357 hPSC line, but it did not affect *RET* expression in hPSCs when compared to the isogenic control line (Figure 5d). Our data suggest that the risk allele *T* in rs2435357 reduces *RET* expression in the early enteric neural progenitors (hNPs), contributing to HSCR pathogenesis.

**Figure 5.**
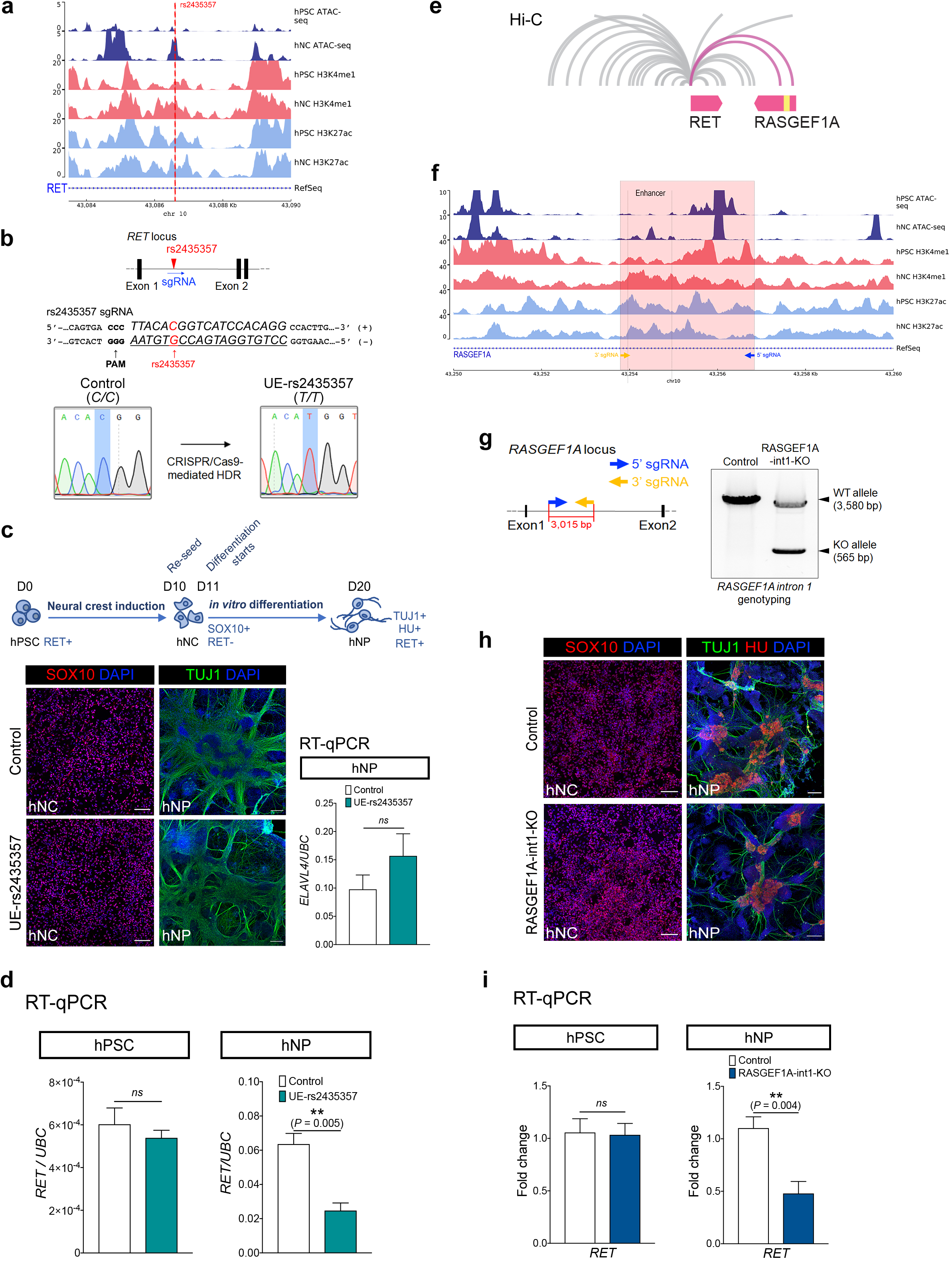
Functional impacts of a HSCR-associated SNP (rs2435357) and the deletion of a novel S-HSCR enhancer on *RET* expression. (**a**) ATAC-seq and ChIP-seq profiles of hPSC and hNC in the intron 1 of *RET* show that rs2435357 is residing in a hNC-specific ATAC-seq peak. (**b**) Location of rs2435357 in the *RET* gene locus and in the sgRNA used for CRISPR/Cas9-mediated HDR for editing the *C* allele to the HSCR-associated risk allele *T*. The electrographs of Sanger sequencing show the successful introduction of the risk allele at rs2435357 in the UE-rs2435357 hPSC line. (**c**) Differentiation strategy to generate human neural crest (hNC) and neuronal progenitor (hNP), and immunostaining of SOX10 and TUJ1 in hNC and hNP of the control and the mutant (UE-rs2435357) lines. Scale bars: (hNC): 100 *μ*m; (hNP): 200 *μ*m. RT-qPCR analysis showing the comparable *ELAVL4* expression level in hNP in the control (*n*=5) and the mutant (UE-rs2435357) (*n*=3) lines. *t-*test, *ns*: not significant. (**d**) RT-qPCR analysis showing *RET* expression in the hPSC and hNP stages of the control (*n*=5) and the mutant (UE-rs2435357) (*n*=3). *t-*test, *ns*: not significant. (**e**) Publicly available Hi-C data from GM12878 cells^84^ are shown using the *RET* locus as anchor. The putative enhancer in intron 1 of *RASGEF1A* is marked in yellow. (**f**) ATAC-seq and ChIP-seq data from hPSC and hNC at the *RASGEF1A* intron 1 locus. (**g**) The design of sgRNAs used for the CRISPR/Cas9 system for deleting the DNA fragment in *RASGEF1A* intron 1. Genotyping reveals the specific deletion of *RASGEF1A* intron 1 in UE-RASGEF1A-int1-KO hPSC line. WT: wildtype; KO: knockout. (**h**) Immunostaining of SOX10, TUJ1 and HU in hNC and hNP of the control and the mutant (UE-rs2435357) lines, respectively. Scale bars: (hNC): 100 *μ*m; (hNP): 200 *μ*m. (**i**) RT-qPCR reveals the expression level of *RET* in the hPSC and hNP stages of the control (*n*=4-5) and the mutant (RASGEF1A-int1-KO) (*n*=6-7). *t-*test, *ns*: not significant.

Next, we studied a novel enhancer (chr1:204456576-204457577) significantly associated with S-HSCR within intron 1 of *RASGEF1A* that is located around 180kbp downstream of the TSS of *RET* (Table 1). This putative enhancer is close to the TSS of *RET* in the three-dimensional (3D) genome architecture in GM12878 cells based on previously published Hi-C data^84^ (Figure 5e). ATAC-seq data also revealed that there was an hNC-specific peak in this region, implying that this enhancer may also be implicated in ENS development (Figure 5f). To investigate the potential effect of this novel enhancer in controlling *RET* expression, we generated a knockout (KO) mutant hPSC line (RASGEF1A-int1-KO) using a CRISPR/Cas9 system^80^ with a pair of specific sgRNAs flanking the target region of intron 1 of *RASGEF1A*. Single hPSC colonies were isolated and genotyped to confirm the deletion of the target region (Figure 5g). After neural induction and neuronal differentiation, we found that the differentiation potential of RASGEF1A-int1-KO hPSCs to make hNCs and hNPs was highly comparable to that of the control. Comparable numbers of SOX10, TUJ1 and ELAVL4 expressing cells were found in the control and the RASGEF1A-int1-KO groups (Figure 5h). However, the deletion significantly reduced *RET* expression in hNPs, but not in hPSCs (Figure 5i), implying that intron 1 of *RASGEF1A* contains a long-range regulator of *RET* in the early enteric progenitors.

### Disruption of NFIA binding in a novel S-HSCR-associated regulatory element in intron 10 of *PIK3C2B* interferes the expression of *PIK3C2B, PPP1R15B* and *SOX13*

We further experimentally studied the functional potential of another S-HSCR associated regulatory element located in intron 10 of *PIK3C2B* (chr1:204456576-204457577). This regulatory element was of particular interest because (1) it contained a strong hNC-associated ATAC-seq peak (Figure 6a); (2) an *A*>*T* variant in this locus (rs551359143) was identified in six of our S-HSCR patients but not in any control subjects; (3) the *A*>*T* conversion was predicted to disrupt the binding motif of NFIA (Table 1), which is a TF with a key role in regulating the differentiation of neural stem cells^44,45,85,86^ (Figure 6a). We generated another mutant hPSC line in which 171bp of the intron 10 of *PIK3C2B* flanking the putative NFIA binding motif was deleted using a CRISPR/Cas9 system. The deletion of the target region was confirmed by genotyping in the PIK3C2B-int10-KO hPSC clone (Figure 6b). The mutant hPSC clone was used to make hNCs and hNPs as described above. Similar to the other enhancers discussed above, the deletion of the fragment did not affect the differentiation potential of the hPSCs to make hNCs and hNPs (Figure 6c). In terms of gene expression, our RT-qPCR revealed a sequential down-regulation of *PIK3C2B* along the enteric neuronal progenitor differentiation (hPSC-to-hNC-hNP). When the 171bp fragment was deleted, the expression of *PIK3C2B* in hNCs and hNPs was significantly increased as compared to the isogenic control, implying that this 171bp fragment probably serves as a negative regulatory element to mediate the sequential down-regulation of *PIK3C2B* along the differentiation path and the loss of it led to aberrant expression of *PIK3C2B* gene in hNCs and hNPs (Figure 6d). We then further demonstrated the involvement of NFIA in mediating *PIK3C2B* expression using luciferase assay. Our data showed that NFIA mainly acts as a repressor and the luciferase activity was significantly up-regulated when the NFIA binding motif was disrupted by introducing the *A*>*T* substitution (Figure 6e). A direct binding of NFIA onto the intron 10 of *PIK3C2B* was demonstrated using gel shift and supershift assay, in which NFIA binding was consistently abolished by the *A*>*T* conversion (Figure 6F). All these results indicate that *PIK3C2B* was negatively regulated by NFIA and the HSCR-associated *A>T* variation would disrupt NFIA binding, leading to the increased expression of *PIK3C2B*.

**Figure 6.**
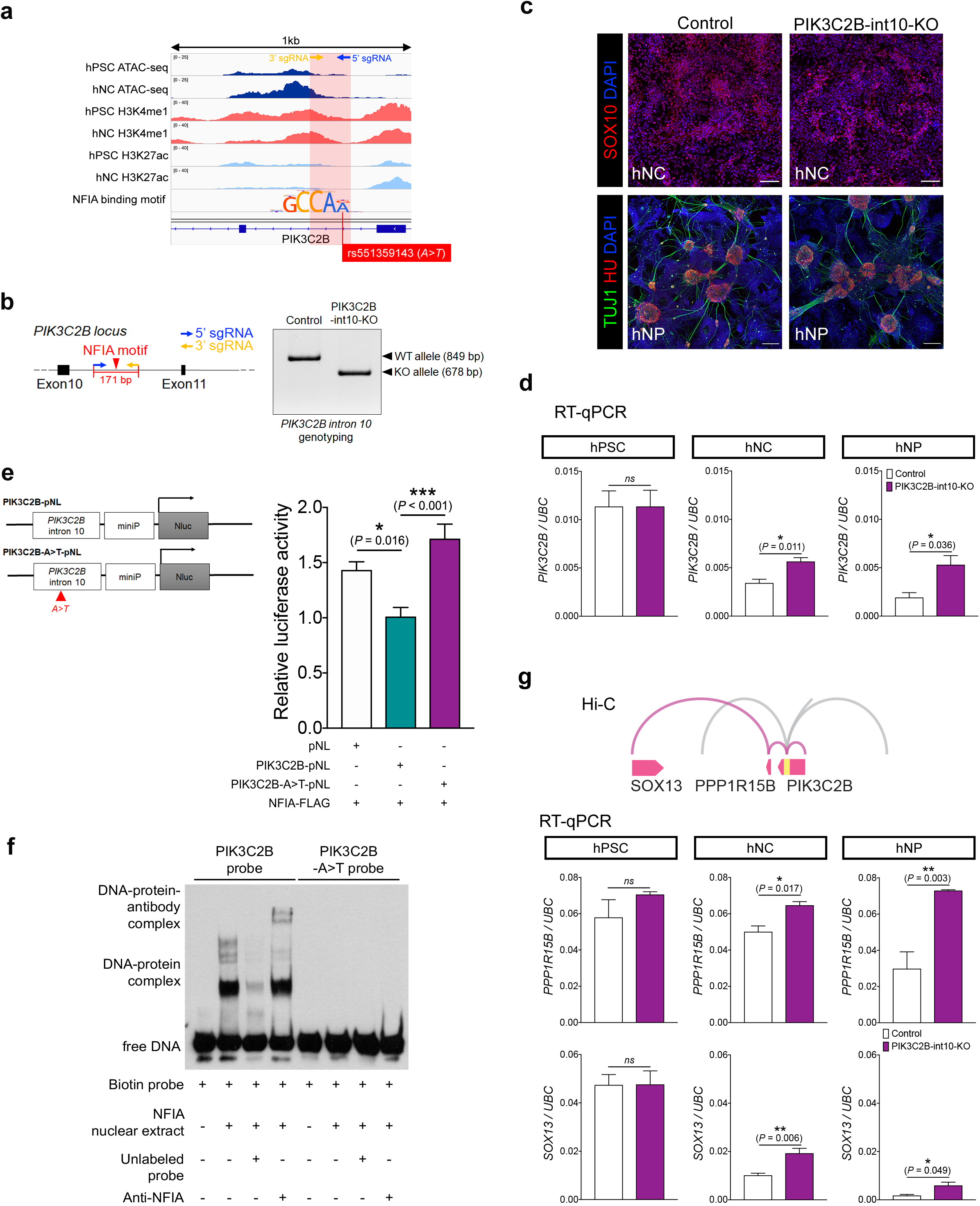
Characterization of a novel S-HSCR-associated regulatory element in the intron 10 of *PIK3C2B*. (**a**) Overview of ATAC-seq and ChIP-seq profiles showing the putative hNC-specific regulatory elements in *PIK3C2B* intron 10 locus. The red shaded region indicates the location of the regulatory element and the line shows the *A*>*T* variant (rs551359143) found exclusively in the S-HSCR cases that disrupts the NFIA binding motif. The motif is not drawn in the same scale as the genomic signal tracks, with magnified characters. (**b**) Design of sgRNAs used for the CRISPR/Cas9 system for deleting the regulatory element. Genotyping reveals the specific deletion of the 171 bp fragment in intron 10 of *PIK3C2B* in the PIK3C2B-int10-KO hPSC line. WT: wildtype; KO: knockout. (**c**) Immunostaining shows that both the control and mutant (PIK3C2B-int1-KO) lines have comparable capability to make hNCs and hNPs. Scale bars: (hNC): 100 *μ*m; (hNP): 200 *μ*m. (**d**) RT-qPCR shows the changes in the expression of *PIK3CB* in different cell stages in the control and the mutant lines. *t-*test, *ns*: not significant. *n*=3-4 per group. (**e**) Design of the constructs used for the luciferase assay. Bar chart indicates the relative luciferase activities when transfected with different sets of constructs as indicated. Three independent assays were performed, each in triplicate. One-way ANOVA. (**f**) Gel mobility shift assays were performed with biotin-labeled probes containing the *PIK3C2B* intron 10 regulatory element with or without the *A*>*T* conversion and the nuclear extract from NFIA-overexpressing cells, in the presence of unlabeled probes or anti-NFIA antibody (0.1 µg). (**g**) Hi-C data from GM12878 cells^84^ at the *PIK3C2B* locus are shown. The putative regulatory element in intron 10 of *PIK3C2B* is marked in yellow. RT-qPCR analysis shows the changes in the expression of *PPP1R15B* and *SOX13* in the control and the mutant lines at different cell stages. *t-*test, *ns*: not significant. *n*=3-4 per group.

Long-range chromatin interactions form topologically associating domains (TAD) and the genes within a TAD are likely to be regulated by common enhancers^15,87^. A previous Hi-C experiment suggests that chromatin loops can be formed between *PIK3C2B* intron 10 and the TSSs of *PPP1R15B* and *SOX13* in GM12878 cells^84^ (Figure 6g). This suggests that the gene expression of *PPP1R15B* and *SOX13* may also be influenced by the S-HSCR-associated *A*>*T* variant in intron 10 of *PIK3C2B*. RT-qPCR revealed that the expression levels of *PPP1R15B* and *SOX13*, similar to that of *PIK3C2B*, were not affected in the hPSC stage but upregulated in the hNC and hNP stages after the deletion of the 171bp fragment (Figure 6g). This result suggests that *PIK3C2B* intron 10 contains an element that can regulate the expression of genes within the same TAD and the usage of this regulatory element may be dynamic during the development of ENS. During neuronal differentiation, this element likely exerts negative control over gene expression as the deletion of it consistently increased gene expression. Interestingly, in the control line, all the genes regulated by this element were downregulated when the cells differentiated into neuronal lineages, suggesting that upregulation of these genes associated with the *A*>*T* variant may impose disease-causing risks during the later stage of enteric NC cell development.

## Discussion

In this work, we have proposed a novel framework, MARVEL, for identifying noncoding genetic variants of potential functional significance. It was designed to overcome various issues in the study of noncoding variants. Specifically, it uses epigenomic data to define a target set of regulatory regions and sequence motifs to estimate the functional impacts of the variants, which help reduce the burden of multiple hypothesis testing and prioritize variants in linkage disequilibrium according to their functional potential. It jointly analyzes different variants within the same regulatory region and offers the option of combining information from different regulatory regions of the same gene, which help capture variants with non-linear interactions and rare signals that converge to the same outcome. The subsequent analysis steps of MARVEL further aggregate the identified genes into functional pathways and investigate the upstream regulators that have their binding sites recurrently perturbed, which provide concrete testable hypotheses for follow-up investigations.

Our application of MARVEL to the WGS data of S-HSCR has clearly demonstrated the advantages of these designs. Starting from 37 millions of noncoding variants observed in the cases and controls, the epigenomic data reduced the number of genomic regions to study to the scale of tens of thousands (of genes and promoters) to hundreds of thousands (of enhancers), and the statistical procedures further selected no more than 10 key motifs from each of these regions among 771 of them. As a result, we were able to obtain some results other than those in the dominating *RET* locus at fairly strong significance levels, which would be impossible in the traditional approach of testing the disease-association of each genetic variant separately since most noncoding variants are rare or have small effect sizes.

Comparing the enhancer-based and promoter-based analyses, we obtained more significant results from the former, showing that it is important to move beyond the immediate TSS-proximal regions when analyzing the noncoding variants. The designs of MARVEL made it possible to study genetic variants in distal enhancers in a computationally and statistically feasible manner.

The gene-based analysis led to findings in chromosomes 9 and 15 not obtained from the enhancer-based or promoter-based analysis. We have found that around these gene loci there were more enhancers with marginally significant associations than other gene loci, suggesting that it was the aggregation of weak signals from multiple regulatory elements that helped identify these gene loci as significantly associated with S-HSCR. The importance of this aggregation idea was further demonstrated in the functional pathway analysis, in which some of the 552 genes loosely associated with S-HSCR were grouped into 5 functional clusters that are highly related to NC migration.

Finally, our analysis of upstream regulators have identified 48 recurrent TFs and they form an extensive network through PPIs. Most interestingly, some of these TFs were also have their corresponding genes identified in the association tests, showing that both the expression of these TFs themselves and their binding at other genes’ regulatory regions are associated with S-HSCR, illustrating two main mechanisms by which genetic variants can exert their functional effects.

Overall, our work has produced a large number of novel regulatory elements, genes and upstream regulators associated with S-HSCR for follow-up investigations. Our genome editing experiments have confirmed that three regulatory elements, including one within the *RET* locus, one within the *RASGEF1A* locus near *RET*, and one within the *PIK3C2B* locus, regulate expression of important genes (*RET, PIK3C2B, PPP1R15B* and *SOX13*) in a cell stage-specific manner. In the first case, we were able to show the effects of a single genetic variant on *RET* expression. In the third case, we experimentally showed that the genetic variant affected the binding of NFIA. A common observation in all three cases is that although expression of some key genes were perturbed by the genome editing, the differentiation potential of the hPSCs into hNCs and hNPs was apparently unaffected. We hypothesize that this was due to the buffering of different regulatory elements and different genes, which causes single perturbations to have little observable effect on cell differentiation and as a result their roles in disease susceptibility and pathogenesis are subtle. In order to have a more complete understanding of the functional importance of the candidate regulatory elements, genes and TFs identified in this study, it would be useful to perform large-scale genome editing assays to systematically perturb them individually and in combination and observe the resulting phenotypes.

## Methods

### Details of the MARVEL framework

#### Required inputs

MARVEL requires two main inputs from the user, namely 1) a list of genetic variants from each subject, which can include single-nucleotide variants and small insertions and deletions, and 2) a set of target regulatory elements. The target regulatory elements required are the enhancers for an enhancer-based analysis, promoters for a promoter-based analysis, and both enhancers and promoters for a gene-based analysis.

#### Reconstruction of sample-specific sequences

For each target regulatory element, the genomic sequence of each subject is reconstructed by merging the supplied genetic variants of this subject into the human reference sequence. Specifically, the reconstructed sequence will contain the variant allele if the variant is either homozygous or heterozygous with one of the two alleles being the reference one. For a heterozygous variant with both alleles different from the reference one, one of them is included in the reconstructed sequence arbitrarily.

#### Motif scores calculation and aggregation

Based on the reconstructed sequence of each target regulatory element, the match (log odds) scores of 717 motifs from the HOCOMOCO^88^ human TF motif database (v11) are computed using MOODS (v1.9.3), with the score set to 0 if the P-value does not pass the default threshold of 10^−5^. When computing these scores, the nucleotide frequency background is taken from all the sequences in the set of target regulatory elements. If a motif has multiple occurrences in a regulatory element, their match scores are added up to give a single score for this motif. For a gene-based analysis, the scores of a motif in different regulatory elements (including both promoters and enhancers) are further aggregated by a weighted sum, where the weight indicates the strength or confidence of each regulatory element in regulating the gene. For example, if high-throughput chromosome conformation capture data are available, the contact frequencies between the promoter of a gene and different enhancers can be used to define the weights of these enhancers, with a stronger weight given for an enhancer with a higher contact frequency with the promoter.

After this step, each target region (enhancer, promoter or gene) has a single motif score profile, which is a vector of 717 motif scores. The motif scores are further standardized such that each motif has a mean score of zero and a standard deviation of one across all the subjects.

#### Identification of important motifs and phenotype-associated target regions

Suppose there are *n* subjects with their phenotype information recorded in the binary vector ***y***, where *y*_*i*_ = 1 if subject *i* is a case and *y*_*i*_ = 0 if subject *i* is a control. For each target region, the standardized score profiles are represented by an *n* × *m* matrix ***X***, where *m* is the number of motifs. Covariates of the *n* subjects are given in an *n* × *v* matrix ***C***, where *v* is the number of covariates.

Based on ***X*** and ***y***, generalized-linear-model-based least angle regression (GLM-LARS^89^) is performed to select a subset of motifs whose match scores in this target region can best explain the phenotypes. We use glmpath^90^ (https://cran.r-project.org/web/packages/glmpath/index.html) to perform GLM-LARS with parameter settings min.lambda=1e-2, max.steps=10, and max.vars=10 such that up to 10 motifs can be selected and the computations are efficient.

To quantify how well the target region’s genetic variations are associated with the phenotypes based on the scores of the selected motifs, we construct two logistic regression models:

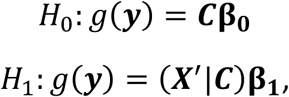

where 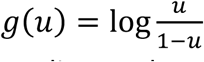 is the logit function, ***X***^′^ is a reduced version of ***X*** retaining only the columns corresponding to the selected motifs, (***X***^′^|***C***) is a matrix that concatenates the columns of ***X***^′^ and ***C***, and **β**_**0**_ and **β**_**1**_ are coefficients. The likelihood ratio between these two models is then used as a test statistic:

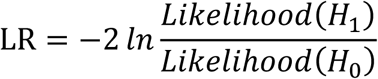

The statistical significance of a target region is evaluated by comparing its LR with those from a permutation set, which involves all the target regions with 100 random permutations of ***y***. For example, when we applied this procedure to perform association tests for the enhancers in the S-HSCR study, since there were 150,828 enhancers in total, each of these LR values were compared against the 15,082,800 values from the permutations to compute a P-value. The P-values of all the target regions are then corrected for multiple hypothesis testing using the Benjamini-Hochberg method^91^.

### Estimation of effect size

We use AUROC to estimate the effect size of each target region. Specifically, after constructing the model *g*(***y***) = (***X***^′^|***C***)**β**_**1**_ for a target region, for each subject, this model is applied to compute a score that indicates how likely this subject has the phenotype. These phenotype scores together with the actual phenotype labels of the subjects are then used to produce the receiver-operator characteristic, and the area under the curve is used as an estimate of the effect size.

### Validation of MARVEL using simulated data

We simulated 3 motif score profiles to evaluate the performance of GLM-LARS in selecting important motifs caused by different types of genetic variants (Supplementary Figure 1). In each scenario, the data for 100 cases and 100 controls were generated.

The first profile (Supplementary Figure 1a) contained 4 motifs (x1-x4) whose match scores were associated with the phenotype due to common genetic variants with moderate effect sizes and 200 motifs (x5-x204) whose match scores were not affected by the genetic variants (for simplicity, here we use the same symbol for both a motif and its score vector). Among the four associated motifs, x1 and x2 were more associated with the phenotype than x3 and x4, and thus the former two were expected to receive larger absolute coefficients in the regression model.

The second profile (Supplementary Figure 1b) contained 4 motifs (x1-x4) whose match scores were associated with the phenotype due to common variants with moderate effect sizes, 1 motif (x5) whose match scores were associated with the phenotype due to less common variants with large effect sizes, 11 motifs (x6-x16) whose match scores were altered in <1% of individuals due to sequencing errors, and 200 motifs (x17-x216) whose match scores were not affected by the genetic variants. In this scenario, x1 and x2 were expected to have the largest coefficients in the regression model, followed by x5, and in turn followed by x3 and x4.

The third profile (Supplementary Figure 1c) was generated using a procedure similar to the one used for the second profile, except that the last motif (x216) was completely correlated with the first motif (x1), with the match score of the former being half of the match score of the latter in every subject. Both x1 and x216 were expected to receive the same non-zero coefficient in the regression model.

To evaluate the statistical testing procedures of the target regions, we simulated 500 motif score profiles. Among them, 10 motif score profiles were associated with the phenotype using the same procedure as the third profile described above. The remaining 490 motif score profiles were generated randomly with 4 motifs whose match scores were sampled from a normal distribution, 12 motifs whose match scores were altered in <1% of individuals due to sequencing errors, and 200 motifs whose match scores were not affected by variants.

### Application of MARVEL to S-HSCR

#### Whole-genome sequencing and variant calling data

In our previous study^8^, WGS was performed on 431 S-HSCR cases and 487 ethnically matched controls. Quality checks and processing of the data were performed, and genetic variants were called from the resulting data using standard methods^8^. We supplied all the identified variants as input to MARVEL, including both common and rare variants.

#### Assay for transposase-Accessible Chromatin with high-throughput sequencing (ATAC-seq)

ATAC-seq was performed as previously described^92^. In brief, around 35,000 FACS-enriched hNC or hPSCs were collected and washed in cold PBS. Transposition reaction was performed according the manufacturer’s protocol for Nextra Tn5 transpoase Nextra kit (Illumina, Cat. No.: FC-121-1030). Transposed DNA fragments were purified by Qiagen MiniElute PCR purification kit (Qiagen). DNA libraries were then prepared by PCR amplification using NEBNext High-Fidelity PCR kit (New England Biolabs) in the presence of barcoded PCR primers (sequences provided in Ref. ^92^). After the PCR amplification, DNA libraries were purified twice by 1.8x AMPure XP beads (Bechman Coulter A63880). The quality of the purified DNA library was assessed by a Bioanalyzer High-Sensitivity DNA Analysis Kit (Agilent). Illumina HiSeq SBS Kit v4 was used for PE101 sequencing (Illumina).

#### Chromatin immunoprecipitation sequencing (ChIP-seq)

2.5-5×10^5^ FACS enriched hNC or hPSC were collected and fixed with 37% formaldehyde for 10 minutes at room temperature. Chromatin was sonicated by Bioruptor Plus UCD-300 (Diagenode, Belgium). ChIP was performed with 5 μg of H3K4me1 or H3K27ac antibodies (Abcam), and normal IgG (inputs as control), respectively, by MAGnify™ Chromatin Immunoprecipitation System (Invitrogen, USA). ChIP-seq libraries were prepared by MicroPlex Library Preparation kit v2 (Diagenode) and Illumina sequencing (Pair-End sequencing of 101bp) were done at The University of Hong Kong, Centre for PanorOmic Sciences (HKU, CPOS).

#### Production and processing of epigenomic data

ChIP-seq targeting H3K4me1 and H3K27ac and ATAC-seq were performed on the hNC and hPSC with two biological replicates for each assay as described above. The raw data were processed using the ENCODE standard ChIP-seq (https://github.com/ENCODE-DCC/chip-seq-pipeline) and ATAC-seq (https://github.com/kundajelab/atac_dnase_pipelines) pipelines, which included read alignment, quality control, reproducibility assessment, and reads pooling. Narrow peaks were then called from the pooled reads using MACS2^93^ with default settings, involving the matched input controls in the case of ChIP-seq.

#### Defining target regulatory regions

ChromHMM^94^ (v1.20) was used to perform genome segmentation based on the ChIP-seq and ATAC-seq peaks. We defined an initial set of enhancers as the genomic segments in chromatin states that emitted both H3K4me1 and H3K27ac marks and overlapped an ATAC-seq peak. These enhancers were then size-normalized to 1kbp each, covering the +/- 500bp regions around the corresponding ATAC-seq peak summits. Target promoters were defined as the +/-500bp regions around all the TSSs of all the genes in GENCODE (v28).

For the negative control study, we collected FANTOM5^25,95^ phase 2 permissive enhancers from all samples, extended the length of each enhancer to 1,000 bp while keeping the enhancer center unchanged, and kept only those having no overlap with the active hNC enhancers defined above.

#### Construction of motif score profiles

For each target enhancer and promoter, we constructed the motif score profile of each subject as described above. For the gene-based analysis, for each gene we considered all its promoters together with all enhancers within 1Mbp from its first TSS. The weight of each regulatory element depends on its genomic distance from the TSS (with the distance of the promoter defined as 0). Specifically, the frequencies of chromatin contact at different distance bins from 0 to 1Mbp were collected from a previous study that performed capture Hi-C on hPSCs^96^. These frequencies were normalized to have a sum of one and these normalized frequencies were used as the weights (Supplementary Table 4).

#### Covariates

In the statistical testing procedure of MARVEL, following previous work^8^, we used the first three principal components of the genetic variant matrix as the covariates.

### Comparing MARVEL with existing single-variant and region-based tests

We compared the enhancer-based results of MARVEL with the results of 3 commonly used association tests, namely the single-variant Wald test and the region-based tests CMC (Combined Multivariate and Collapsing)^97^ and SKAT-O (Optimized Sequencing Kernel Association)^31^. All three tests were performed using RVTESTS^98^ with the same covariates as MARVEL. Following the common practice^99–101^, Wald tests were performed on biallelic common variants with minor allele frequency (MAF) larger than 0.01, while the 2 other tests were conducted on rare variants (MAF<0.01). For both cases, only the variants within the same set of hNC enhancers used by the MARVEL enhancer-based analysis were considered. The P-values from each testing approach were separately corrected for multiple hypothesis testing using the Benjamini-Hochberg method. Motif scanning was performed on +/-20bp around each variant using MOODS based on the same motif set, P-value threshold and nucleotide background frequencies as in the MARVEL enhancer-based test.

For the loosely associated variants identified by the Wald test, pairwise r^2^ values were computed using PLINK (v1.9) with the ‘--ld’ parameter^102,103^. The variant pairs with an r^2^ value higher than 0.9 were defined to be in linkage disequilibrium.

### Analysis of functional pathways

The 552 genes used in the study of functional pathways were the union of the genes loosely associated with S-HSCR from the enhancer-based, promoter-based and gene-based analysis results. For the enhancer-based results, the loosely associated genes included all genes within 100kbp from each loosely associated enhancer, or the gene closest to it when there were none.

The functional interactions were obtained from Reactome^60^ (v7.2.0) (https://reactome.org/tools/reactome-fiviz), which contained manually curated functional interactions among over 60% of human proteins. The Reactome-Flviz plugin of Cytoscape^60^ was used to obtain and visualize the functional interactions.

To evaluate the functional connectedness of the 552 genes, we counted the number of them having at least one functional interaction with another gene in this set and the total number of interactions among them. We then repeated this same procedure for 1,000 random sets of 552 genes, and computed the P-value as the number of random sets having a larger number of connected genes/interactions than the 552 genes loosely associated with S-HSCR.

When exploring the functions of the 552 genes, an expanded gene set was created by first adding 26 known HSCR genes, including *BACE2, DNMT3B, ECE1, EDN3, EDNRB, FAT3, GDNF, GFRA1, KIAA1279, L1CAM, NRG1, NRG3, NRTN, NTF3, NTRK3, PHOX2B, PROK1, PROKR1, PROKR2, PSPN, SEMA3A/C/D, SOX10, TCF4*, and *ZFHX1B* ^8–13^. After that, genes involved in ENS function or NC migration that had Reactome functional interactions with at least one gene already in the expanded gene set were further added to it, including *CDC42*^61^, *CUL1*^47^, *ERBB2/3/4*^61^, *GNAI1*^104^, *GRB2*^105,106^, *NRP1/2*^61^, *PAX3*^71,107^, *PLXNB1*^108^, *ROBO1/2/3*^61^, *SLIT1/2/3*^61^, and *TCF12*^107^.

### Analysis of mouse trunk NC scRNA-seq data

Processed mouse trunk NC scRNA-seq data^70^ were downloaded from http://pklab.med.harvard.edu/ruslan/neural.crest.html. The expression profiles were extracted from the ‘wgm2’ data matrix, which had been batch-adjusted and mean-variance normalized as described in the original paper^70^. The crestree (https://github.com/hms-dbmi/crestree) R package was used to visualize the expression profiles of Myc and its binding targets across NC lineages. In-house R scripts and the pheatmap library (https://cran.r-project.org/web/packages/pheatmap/pheatmap.pdf) were used to produce the expression heatmaps. To select genes with stage-specific expression for visualization, principal component analysis was performed on the expression matrix with the genes treated as features. The 10 genes with the largest absolute loading in each of the top three principal components were included in the visualization.

### Analysis of recurrent TFs

To identify the recurrent TFs, all motifs selected by GLM-LARS from the full set of 200 enhancers loosely associated with S-HSCR were collected. 10,000 random sets of 200 enhancers were then formed by sampling from all enhancers, and GLM-LARS was applied to each of these sets. For each motif, its total model coefficients across all enhancers in a set was used as a test statistic, and a P-value was computed by comparing the test statistic obtained from the loosely associated enhancers with those from the random enhancer sets, to see if the former is significantly larger than the latter. These raw P-values were then corrected by the Benjamini-Hochberg method to control the FDR at 0.05.

The PPIs among the recurrent TFs were obtained from STRING^109^ (v11). We included only the interactions based on manual curation or experimental evidence with a combined score >0.4.

### Functional studies

#### Cell culture

A control hPSC line (UE02302) was established as previously described^110^. hPSCs were maintained in Matrigel (Corning)-coated plate in mTeSR1 medium (Stem Cell Technologies) in a 37 °C humidified 5% CO2 incubator. The hPSCs were regularly passaged by treating with Dispase (Stem Cell Technologies).

Neural crest induction was performed according to a previously described protocol^63^. In brief, hPSCs were dissociated into single cell suspension by Accutase (Millipore) and plated on Matrigel-coated plate in a density of 5×10^4^ cells cm^-2^ in ES cell medium containing 10 ng/mL fibroblast growth factor 2 (FGF2, Peprotech). The differentiation was started by replacing ES cell medium with KSR medium and gradually switched to N2 medium from day 4 to day 10. To differentiate hPSCs to hNC cells, the cells were treated with 100 nM LDN193189 (Stemgent) from day 0 to day 3, 10 μM SB431542 (Abcam) from day 0 to day 4, 3 *μ*M CHIR99021 (Stemgent) from day 2 to day 10 and 1 *μ*M retinoic acid from day 6 to day 10.

For neuronal differentiation of hNCs to hNPs, hNC cells were dissociated into single cell suspension by Accutase (Millipore) at day 10. For the study of the regulatory element in *PIK3C2B* intron 10, the harvested cells were pelleted and resuspended with N2 medium containing 10 ng/mL FGF2 and 3 *μ*M CHIR99021in a density of 5×10^3^ cells *μ*l^-1^. 2.5 ×10^4^ hNC cells were seeded as droplets on polyornithine/laminin/fibronectin-coated surface. For the *RET*-associated study, the harvested hNC cells were subjected to fluorescence-activated cell sorting (FACS) and hNC cells which were positive to both HNK-1 (BD Biosciences #560845) and p75^NTR^ (Miltenyi Biotec #130-091-917) were sorted by BD FACSAria III Cell Sorter. 5 ×10^4^ sorted cells were seeded as droplets on polyornithine/laminin/fibronectin-coated surface. Neuronal differentiation was initiated by replacing the medium with N2 medium containing 10 ng/mL BDNF (Peprotech), 10 ng/mL GDNF (Peprotech), 10 ng/mL NT-3 (Peprotech), 10 ng/mL NGF (Peprotech), 1 *μ*M cAMP (Sigma) and 200 *μ*M ascorbic acid (Sigma). The hNC cells differentiated into hNP cells in 9 days.

#### Plasmid constructions

Human codon-optimized high fidelity Cas9 nuclease construct with GFP tag (pSpCas9(BB)-2A-GFP (PX458))^80^ was obtained from Addgene (#48138). Two sgRNAs were designed using CRISPR-Cas9 guide RNA design checker from Integrated DNA Technologies. Oligos for sgRNA cloning are listed in

Supplementary Table ***5***. The annealed sgRNA oligos were ligated with *Bbs*I-linearized Cas9 construct using Blunt/TA ligation mix (New England Biolabs). For the sgRNA targeting the rs2435357 locus, the annealed sgRNA oligo was ligated with *Afl*II-linearized gRNA cloning vector^111^ (Addgene #41824) using Blunt/TA ligation mix.

For luciferase assay, *PIK3C2B* intron 10 fragment was amplified from the control hPSC genomic DNA and *NFIA* ORF was amplified from the control hNP cDNA using Q5 Hot Start High-Fidelity DNA Polymerase (New England Biolabs). *PIK3C2B* intron 10 fragment was cloned into NanoLuc luciferase reporter construct (pNL3.2[*NlucP/minP*]) (Promega #N1041) to generate PIK3C2B-pNL construct while *NFIA* ORF was cloned into pFLAG-CMV expression plasmid to generate NFIA-FLAG construct. The *A*>*T* variant was introduced to PIK3C2B-pNL construct by site-directed mutagenesis using QuickChange Lightning Site-Directed Mutagenesis Kit (Agilent) to generate PIK3C2B-*A*>*T*-pNL construct. The cloning primers and mutagenesis primers are listed in Supplementary Table 5.

#### Generation of new hPSC lines using CRISPR/Cas9 system

For the generation of UE-rs2435357 hPSC line, 1×10^6^ UE control hPSCs were transfected with 2 µg sgRNA construct, 20 µg ssODNs and 4 µg pSpCas9(BB)-2A-GFP construct using Human Stem Cell Nucleofector Kit 2 (Lonza). For the generation of UE-RASGEF1A-int1-KO and PIK3C2B-int10-KO hPSC lines, 2×10^5^ UE control hPSCs were transfected with a pair of pSpCas9(BB)-2A-GFP constructs containing the specific sgRNAs (350 ng per construct) using P3 Primary Cell 4D-Nuclecfector X Kit (Lonza). The transfected cells were plated in Matrigel-coated dish and cultured for 2 days. hPSCs expressing GFP were sorted as single cells into Matrigel-coated 96-well plate with BD FACSAria III Cell Sorter. The sorted cells were expanded for 2 weeks and genotyped to confirm the site-specific conversion or the deletion of the target regions.

### Quantitative RT-PCR (RT-qPCR)

Total RNA from hPSCs, hNCs and hNPs was extracted by RNeasy Mini Kit (Qiagen). RNA concentration was determined by Nanodrop 1000 (Thermo Fisher Scientific) and 100ng or 500 ng RNA was then reverse-transcribed to cDNA using HiScript II Q RT SuperMix (Vazyme). The expression levels of the target genes were quantitated using real-time quantitative RT-PCR or Droplet digital PCR (ddPCR). For real-time quantitative RT-PCR, diluted cDNA samples were amplified by Luna Universal Probe qPCR Master Mix (New England Biolabs) using specific TaqMan Gene Expression Assay (*SOX13*, Assay ID: Hs00232193_m1; *ELAVL4*, Assay ID: Hs00956610_mH; *PIK3C2B*, Assay ID: Hs00898499_m1; *PPP1R15B*, Assay ID: Hs03044848_m1; *RET*, Assay ID: Hs01120027_m1; *UBC*, Assay ID: Hs00824723_m1; *18S*, Assay ID: Hs99999901_s1) (Thermo Fisher Scientific) with PCR profiles of 95 °C (1 min) followed by 45 cycles of 95 °C (15 s) and 60 °C (30 s). Fluorescence was measured by ViiA 7 Real-Time PCR System (Thermo Fisher Scientific) at the end of each cycle. Droplet digital PCR (ddPCR) was used to measure *RET* expression in hNPs. 1 µl cDNA samples were mixed with ddPCR Supermix for Probes (Bio-rad #186-3010) and TaqMan Gene Expression Assay probes of *RET* and *UBC* (Thermo Fisher Scientific). The reaction mixtures were then loaded into the sample wells of DG8 Cartridge (Bio-rad #186-4008), followed by 70 µl of Droplet Generation Oil for Probes (Bio-rad #186-3005) into the oil wells. The cartridge was then placed into QX200 Droplet Generator (Bio-rad) for droplet generation. After droplet generation, the reaction droplets were transferred into a 96-well plate and sealed with foil PCR plate heat seal (Bio-rad #181-4040) and proceeded to thermal cycling with profiles of 95 °C (10 min) followed by 40 cycles of 95 °C (30 s) and 60 °C (1 min) and then deactivation at 98 °C for 10 min. The end-point fluorescence signals from the reaction droplets were then measured by QX200 Droplet Reader (Bio-rad). Each individual sample was assayed in triplicate and gene expression was normalized with *UBC* or *18S* expression.

### Gel shift assay

3×10^5^ HeLa cells were seeded to each well of 6-well plates and cultured in DMEM supplemented with 10% fetal bovine serum and 1% penicillin/streptomycin (Life Technologies) 24 hours before transfection. 2 µg NFIA expression construct (NFIA-FLAG) was transfected to each well of cells by FuGENE HD Transfection Reagent (Promega). Nuclear extracts containing the NFIA protein were extracted from 3 wells of transfected cells using nuclear and cytoplasmic extraction kit (Thermo Scientific). 1 mM ssODNs (PIK3C2B: 5’-CGC AAG AGC TCT TCA GAA ATG GAT GCC AAG TGT GTC TCC TCT TCC TGA-3’ and PIK3C2B-*A*>*T*: 5’-CGC

AAG AGC TCT TCA GAA ATG GAT GCC ATG TGT GTC TCC TCT TCC TGA-3’) derived from the intron 10 of *PIK3C2B* were biotin-labeled using Biotin 3’-End DNA Labeling Kit (Thermo Scientific). Biotin-labeled ssODNs were then annealed with reverse complimentary ssODNs to generate biotin-labeled probes. Gel shift assay was performed by mixing the nuclear extracts with biotin-labeled probes according to the manufacture’s protocol (LightShift Chemiluminescent EMSA Kit; Thermo Scientific). In brief, 20 fmol biotin-labeled probes were mixed with 1 µg NFIA nuclear extract in 1X binding buffer containing 50 ng/µl Poly (dI.dC), 0.05% NP-40, 6% glycerol, 60 mM KCl, 1 mM EDTA and 5 mM MgCl2. The binding reactions were incubated for 20 minutes at room temperature. For competition assays, 4 pmol unlabeled probes were added to the mixture before adding the biotin probes. For supershift assays, 0.1 µg anti-NFIA (Sigma, HPA006111) were added to the mixture in the final step before incubation. The binding reactions were resolved in 5% nondenaturing TBE pre-cast gel (Bio-rad) using Mini-PROTEAN® Electrophoresis System (Bio-rad) and then transferred to Biodyne B Nylon Membrane (Thermo Scientific) in 0.5X TBE buffer. After cross-linking, biotin-labeled probes on the membrane were detected using Chemiluminescent Nucleic Acid Detection Module.

### Luciferase assay

1.5 ×10^5^ SH-SY5Y cells were seeded to each well of 24-well plates and cultured in 1:1 MEM:F-12 mix supplemented with 10% fetal bovine serum, 1% non-essential amino acids, 1% sodium pyruvate and 1% penicillin/streptomycin (Life Technologies) 24 hours before transfection. 50 ng control firefly luciferase construct (pGL3-control), 150 ng NanoLuc luciferase constructs (pNL3.2 or PIK3C2B-pNL or PIK3C2B-*A*>*T*-pNL) and 25ng NFIA expression construct (NFIA-FLAG) were transfected into the cells using jetPRIME transfection reagent (Polyplus Transfection) according to the manufacturer’s protocol. Luciferase activities were detected with Nano-Glo Dual-Luciferase Reporter Assay System (Promega) and measured by VICTOR Nivo Microplate Reader (PerkinElmer).

### Immunostaining

hNC cells and hNP-D9 cells were fixed in 4% PFA in PBS for 20 min at room temperature. Fixed cells were blocked with blocking solution (1% BSA, 0.1% Triton X-100 in PBS) at room temperature for 1 h. The blocked cells were then incubated with primary antibodies (mouse anti-SOX10 (1:500, R&D Systems, MAB2864), rabbit anti-TUJ1 (1:1000, Abcam, ab18207) and mouse anti-HU (1:1000, Life Technologies, #A-21271) at 4 °C for 1 overnight. After washing, the sections were incubated with respective secondary antibodies at room temperature for 2 h. Secondary antibodies were conjugated with Alexa Fluor 488/594 (Life Technologies, 1:500). After washing, the stained cells were mounted with ProLong Diamond Antifade Mountant with DAPI (Life Technologies). Fluorescence images were acquired by Carl Zeiss LSM780 or LSM800 confocal microscope.

## Code availability

The source code and compiled programs of MARVEL are available at https://github.com/fuxialexander/marvel.

## Acknowledgements

ATAC-seq, ChIP-seq, RNA-seq were performed in Center for PanorOmic Sciences, The University of Hong Kong. Confocal imaging were performed with equipment maintained by Li Ka Shing Faculty of Medicine Faculty Core Facility.

This project is supported by Hong Kong Research Grants Council Theme-based Research Scheme T12C-714/14-R. K.Y.Y. is additionally supported by Hong Kong Research Grants Council Collaborative Research Funds C4045-18WF, C4054-16G, C4057-18EF, and C7044-19G and General Research Funds 14170217 and 14203119, the Hong Kong Epigenomics Project (EpiHK), and the Chinese University of Hong Kong Young Researcher Award and Outstanding Fellowship. This work described in this paper is substantially supported by a HMRF grant (Project no.: 06173306) and a General Research Fund (HKU 17108019) from the Department of Health and the Research Grants Council of Hong Kong Special Administrative Region, China Hong Kong, respectively to E.S.W.N. K.N.C.L. is supported by the Hong Kong PhD Fellowship from the Research Grants Council of Hong Kong.

## Contributions

P.K.H.T., E.S.W.N. and K.Y.Y. conceived this study. A.X.F. and K.Y.Y. developed the MARVEL framework. A.X.F., C.S.M.T., M.M.G.B., P.C.S. and K.Y.Y. analyzed the sequencing data. R.N., F.P.L.L. S.T.L. and E.S.W.N. performed the epigenomics experiments. S.T.L. and Z.L. performed RNA-seq analysis of hNC. A.X.F. and K.Y.Y. applied the MAVEL framework on the S-HSCR data. K.N.C.L. and E.S.W.N. performed the functional experiments and analyzed the data. A.X.F., K.N.C.L., E.S.W.N. and K.Y.Y. prepared the manuscript. P.C.S., P.K.H.T., E.S.W.N. and K.Y.Y. supervised the project.

